# Short-Duration Fluid Shear Stress Enhances LPS-Driven Macrophage Pro-Inflammatory Response in a Magnitude-Dependent Manner

**DOI:** 10.64898/2026.11.16.766992

**Authors:** Parand Shokrani, Rachael Dillon, Hannah Prendeville, Eimear B Dolan

## Abstract

Macrophages are mechanosensitive cells that integrate physical and biochemical cues to shape inflammatory responses. Mechanical stimuli such as stretch, strain, and matrix stiffness have been extensively studied; however, the effects of fluid shear stress (FSS) on macrophage function remain relatively poorly understood. In this study, we investigated how mild-magnitude, short-duration FSS applied alone or in combination with Lipopolysaccharide (LPS), affects bone marrow-derived macrophages (BMDM) phenotype and function. FSS alone had no effect on naïve macrophages, and lower magnitude shear stress did not produce any significant effects under any condition. However, when macrophages were exposed to 0.8 Pa FSS prior to LPS stimulation, tumour necrosis factor (TNF) secretion was significantly increased, and IL-10 production decreased compared with static controls. This magnitude-dependent priming effect was also sequence-specific, as FSS applied after LPS stimulation did not produce any significant changes. These findings demonstrate that mild FSS can mechanically prime macrophages to amplify the inflammatory response but only above a threshold magnitude and when applied before biochemical stimulation. Our results highlight the importance of both shear magnitude and stimulation order in determining macrophage mechanosensitivity, with implications for understanding immune cell behaviour in mechanically dynamic microenvironments and how inflammation can be affected by the mechanical forces *in vivo*.

## Introduction

Macrophages are highly plastic innate immune cells that play a key role in host defence, inflammation, repair, and maintaining tissue homeostasis[1]. Their activation state is regulated not only by cytokines and pathogen-associated signals, but also by physical cues in their environment, such as fluid shear stress (FSS), stretch, and matrix stiffness, which can influence inflammatory signalling and cytokine production[2,3].

Cells encounter a broad range of mechanical cues throughout their lifespan, including FSS in vasculature[4], cyclic stretch and pressure in mechanically active organs such as the lung[5,6] and heart[7], as well as altered matrix stiffness in fibrotic and tumour microenvironments[8,9]. Monocytes are exposed to shear stress while circulating in the blood and during adhesion to the vascular endothelium[10]. Once resident within tissues, macrophages no longer experience vascular flow as such but can instead encounter FSS generated by interstitial fluid movement[11]. Macrophages express a myriad of mechanosensitive receptors and ion channels, including integrins such as αMβ2 (CD11b)[12] as well as mechanosensitive ion channels such as PIEZO1[13] and transient receptor potential vanilloid 4 (TRPV4)[14], enabling them to respond to mechanical forces in their microenvironment. External mechanical forces are sensed by these mechanoreceptors, which convert these mechanical signals into intracellular signalling cascades that affect cell proliferation, differentiation, migration, and gene expression[10,13–15].

Mechanical cues can be particularly relevant during infection and acute inflammation. For instance, increased microvascular permeability can lead to tissue oedema, fluid accumulation, and elevated interstitial fluid flow, creating a physiologically relevant mechanical cue that macrophages can sense and respond to[16]. Consistent with this, interstitial flow has been shown in 3D tissue models to promote macrophage migration and support a phenotypic switch by upregulating Arginase-1 (ARG1) and transforming growth factor beta (TGF-β) through β1-integrin/Src-dependent activation of STAT3/6, highlighting the potential of FSS in directing macrophage function *in vivo*[11]. However, the response to mechanical stimulation is context-dependent and may vary with the type, magnitude, and exposure duration of the mechanical force, cell type, and concurrent biochemical signals[17–19].

Although macrophage mechanobiology has been extensively explored in relation to matrix stiffness[20–22], cyclic stretch[23–25], and hydrostatic pressure[26,27], the effects of FSS on macrophage responses remain comparatively less well understood. For example, Jui et al. showed that subjecting human monocyte-derived macrophages (MDMs) to shear stress at 15 and 35 dynes/cm^2^ for 3, 24, or 48 hr (via a cone-and-plate viscometer) can drive a pro-inflammatory response. RT-qPCR results showed an upregulation of tumour necrosis factor (TNF) at 24-48 hr, with downregulation of TGF-β1, specifically observed at 24 hr in the 35 dynes/cm^2^ group[28]. On the other hand, interstitial flow (∼3 µm/s), applied using a 3D microfluidic and trans well flow chamber system, has been shown to drive bone marrow-derived macrophages (BMDMs) toward an anti-inflammatory phenotype through upregulation of ARG-1, TGF-β, CD206, CD163 and transglutaminase 2 (TGM2) markers, measured by Western blot, and CD163 and CD206 measured through immunofluorescence[11]. These contrasting findings may reflect differences in FSS magnitude, exposure duration, flow pattern, culture environment, and the presence of biochemical stimulation alongside mechanical cues. Therefore, there remain many critical gaps in our understanding of how FSS affects macrophage inflammatory state. Studies examining the interaction between biochemical polarisation signals (e.g., Lipopolysaccharide (LPS)/Interferon-gamma (IFN-γ)) and concurrent mechanical stimulation have been limited, while such combinatorial stimulation is physiologically relevant[19]. Additionally, several studies investigating the effects of FSS on macrophages have applied shear stress for long durations (up to 48 hr)[18,28,29], suggesting that relatively sustained exposure is often tested to elicit measurable changes in macrophage phenotype and inflammatory signalling. While longer FSS exposure durations offer valuable insight into sustained mechanotransduction, prolonged exposure to constant, unidirectional shear *in vitro* may not fully recapitulate the mechanical environment experienced by tissue macrophages *in vivo*, where interstitial fluid flow is generally low and can vary spatially and temporally[30]. During vascular recruitment, adhesion, and initial contact with endothelial or biomaterial surfaces, cells are transiently and dynamically exposed to FSS[31]. Monocyte-endothelial binding events under flow can occur on sub-second timescales, with contact durations as short as 0.2 s contributing to adhesion probability, and adherent leukocytes can respond mechanically to an imposed shear step within seconds by recoiling and stiffening[32,33]. These rapid mechanical responses suggest that mechanosensing can occur on timescales substantially shorter than those commonly used to assess downstream macrophage phenotypic changes. Therefore, experimental models with shorter shear stress exposure may better capture early *in vivo* mechanobiological responses than prolonged regimens[34].

This study investigates how short-duration FSS, alone and in combination with LPS, affects macrophage phenotype and function. This has implications for understanding macrophage behaviour at sites where mechanical forces and inflammation coincide, such as medical implants, where local inflammation at the implant-tissue interface can act alongside local mechanical forces to jointly drive monocyte/macrophage activation and shape the foreign body response[19]. This approach will clarify how short-term mechanical stimulation integrates with biochemical activation to shape macrophage responses, providing insight into how these cues may interact *in vivo*.

## Materials and methods

### Bone-Marrow derived macrophage isolation

Bone marrow was harvested from the tibia and femur bones from 6–8-week-old female C57BL/6 mice housed at the University of Galway. All procedures were approved by the University of Galway Animal Care Research Ethics Committee (ACREC) and conducted under a project authorisation granted by the Animal Care Research Ethics Committee at the University of Galway (project authorisation No. 2024.01.001Amend 2501), in accordance with European Union Directive 2010/63/EU and Irish S.I. No. 543/2012 (European Union (Protection of Animals used for Scientific Purposes) Regulations 2012). Red blood cells were lysed, and cells were resuspended in Dulbecco’s Modified Eagle Medium (DMEM) containing 10% foetal bovine serum (FBS), 1% penicillin-streptomycin (P/S), and 20% L929-conditioned medium (LCM). 1×10^6^ cells/mL were plated in tissue culture-treated petri dishes, and macrophages were differentiated over 7 days.

### Preparation of L929-conditioned medium

L929 fibroblast cells (NCTC clone 929 ATCC-CCL-1) were cultured in DMEM containing 10% FBS and 1% P/S, at a density of 1×10^6^ cells/mL for 8 days. The LCM was collected, filtered through 0.2 µm filters, and stored at –20C. This conditioned medium is rich in macrophage colony-stimulating factor (M-CSF), which promotes differentiation of bone-marrow precursor cells into mature macrophages[35].

### Activation of BMDMs

Following differentiation, BMDMs were harvested and plated on Epredia™ SuperFrost Plus™ Adhesion glass slides at a density of 3.5×10^5^ cells/cm^2^ in DMEM containing 10% FBS, 1% P/S, and 5% LCM. Cells were left to adhere overnight and then stimulated with 10 or 100 ng/mL lipopolysaccharide (LPS) from Escherichia coli O111:B4 (Merck L2630) for 24 hr.

### Fluid Shear Stress (FSS) Exposure

Fluid shear stress exposure was performed using a custom-designed parallel plate system[36]. Fig. 1A shows a schematic summary of the experimental setup. The setup consisted of a syringe pump (World Precision Instruments AL-1600), parallel-plate chambers, inlet and outlet tubing, stopcocks for media inlet and outlet, gas outlet, and a waste collection reservoir. The cell-seeded slides containing the cell monolayer were placed into the slot at the bottom of the flow chamber, which had a length of 140 mm, a width of 18 mm, and a height of 0.2 mm.

**Figure 1:**
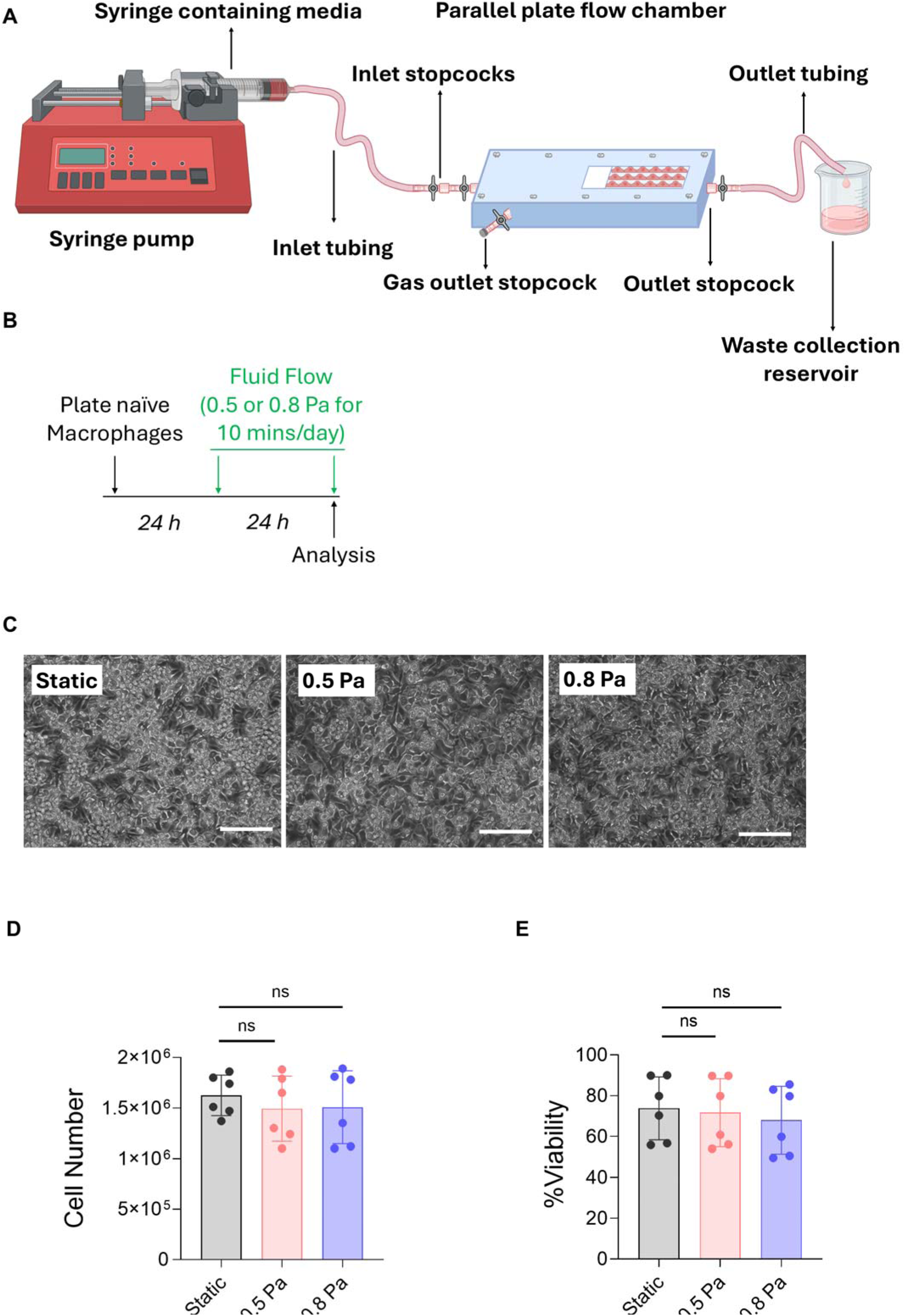
A) Schematic of the parallel plate flow chamber setup and its components. B) Timeline of the study, including exposure of naïve BMDMs to FSS (0.5 or 0.8 Pa, for 10 min/day at 0.5 Hz for two rounds, 24 hr apart), and static controls; C) Microscopic images of the naïve BMDMs (scale bar: 100 μm), D) cell numbers measured using a Luna cell counter and E) percentage of viable cells measured using Live/Dead Aqua staining and flow cytometry. Data are from n = 6 slides from N = 2 independent experiments. One-way ANOVA with Dunnett’s correction was used, and data are represented as mean ± Standard deviation.

Serum-free DMEM culture medium was circulated using a peristaltic syringe pump to create laminar flow at flow rates of 4 mL/min and 6.4 mL/min, inducing FSS of 0.5 and 0.8 Pa, respectively. Cells were subjected to two rounds of FSS for 10 min at 0.5 Hz (1 second pumping, 1 second pause), once every 24 hr. Wall shear stress was calculated using Equation (1) for steady, fully developed laminar flow:

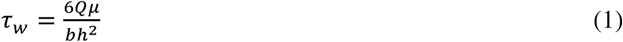

where Q is the flow rate, μ is the fluid viscosity, and b and h are the width and height of the parallel plate flow chamber, respectively[36,37].

For static controls, the slides were placed in the parallel-plate flow chamber with the serum-free media on top, but the media was not circulated through the chamber.

### Microscopy Imaging

Leica DMi1 inverted microscope was used to take microscopic images. Images were taken directly after the second fluid flow exposure at 200X magnification. The scale bars (100 μm) are indicated in the images.

### Flow cytometry

BMDMs were first stained with Live/Dead Aqua viability dye prepared in PBS and incubated in the dark at room temperature for 20 min. Cells were then washed with PBS, and a master mix containing extracellular fluorochrome-labelled antibodies prepared in FACS buffer (PBS containing 2% FBS) was added (Supplementary Table 1). Cells were incubated in the dark at room temperature for 30 min. Cells were then washed with FACS buffer and fixed using the IC fixation buffer (Invitrogen, 12730010). Cells were acquired on the BD FACS Canto II^TM^ flow cytometer, and data were analysed using FlowJo V10 software.

Cells were gated by size (Forward Scatter Area (FSC-A)) and complexity (Side Scatter Area (SSC-A)), and doublets were removed (FSC-A, FSC-H); then live cells were gated from the single-cell population. Macrophages were then gated based on CD11b and F4/80 co-expression. Further analyses were performed on CD11b^+^F4/80^+^ cells.

### Enzyme-Linked Immunosorbent Assay (ELISA)

The concentrations of TNF, IL-6, and IL-10 in cell supernatants were measured using ELISA MAX^TM^ Standard kits (BioLegend), according to the manufacturer’s instructions. Optical density values were measured at 450 nm using a BMG FluoStar OMEGA plate reader. Cytokine concentrations were calculated using a 4-parameter fit curve.

### Statistical Analysis

One-way ANOVA with Dunnett’s correction was used to assess cell viability, cell number, marker expression, and cytokine production between groups. All data are presented as mean ± standard deviation. GraphPad Prism 11.0.0 was used for all statistical analyses. The number of replicates is indicated in the figure legends. BMDMs were seeded onto three slides per experimental repeat and exposed to fluid shear stress or maintained under static culture as a control. Each experiment was independently repeated twice (N = 2), with three slides processed in parallel within each repeat, yielding six slides per condition in total (n = 6). Statistical comparisons were performed at the slide level, with each FSS condition compared to the static control. (Statistical significance was defined based on *p ≤ 0.05, **p ≤ 0.01, ***p ≤ 0.001).

## Results

### Short-duration FSS does not induce functional polarisation of naïve BMDMs

To investigate the effects of FSS on naïve macrophages, naïve BMDMs adhered to glass slides were exposed to FSS using a custom-designed parallel plate system (Fig. 1A). Cells were subjected to two rounds of FSS at 0.5 or 0.8 Pa, for 10 mins at 0.5 Hz, once every 24 hr. As a control, cells were placed in the parallel-plate system but not exposed to FSS (static control; study timeline in Fig. 1B).

FSS at 0.5 or 0.8 Pa did not affect cell adherence to the glass slides, and there were no differences in macrophage numbers between the static and FSS groups (Fig. 1C-D). Microscopic images demonstrate that BMDMs retain an elongated, spindle-like morphology following FSS at 0.5 or 0.8 Pa, typical of differentiated, adherent cells (Fig. 1C). Furthermore, FSS did not affect macrophage survival, as cell viability was similar across static and FSS groups (Fig 1E).

In naïve BMDMs, FSS did not alter the percentage of cells expressing CD11b (integrin expressed by macrophages), CD86 (marker of classical activation) or CD206 (marker of alternative activation) (Fig. 2A-C). Furthermore, expression levels of CD11b, CD86, or CD206 as measured by mean fluorescence intensity (MFI) were not altered by FSS (Fig. 2G-I). TNF, IL-6, and IL-10 cytokines were not detected in the cell supernatants in any of the groups, which is unsurprising as the cells were not stimulated with any biochemical ligands in this experiment (Fig. 2J-L).

**Figure 2:**
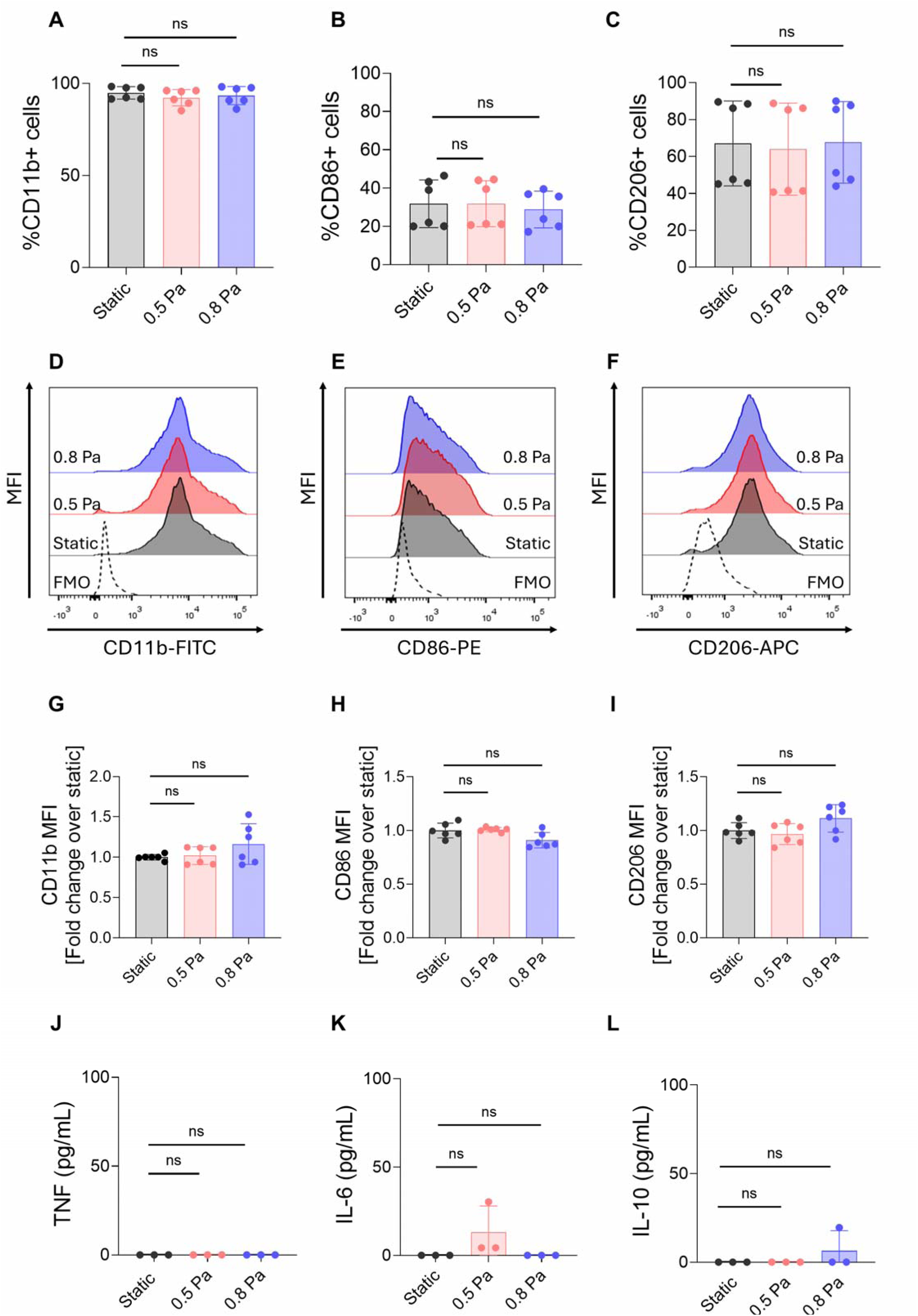
Naïve BMDMs seeded on glass slides and exposed to FSS (0.5 or 0.8 Pa, for 10 min/day at 0.5 Hz for two rounds, 24 hr apart). Percentage of cells expressing each marker, including CD11b (A), CD86 (B), and CD206(C); representative histograms depict mean fluorescence intensity (MFI) of CD11b, CD86, and CD206 in naïve BMDMs exposed to FSS (D-F). Pooled data show that FSS does not affect CD11b, CD86 or CD206 expression (G-I), and cytokine production in the supernatant measured by ELISA (J-L). Data are from n = 6 slides from N = 2 independent experiments for flow cytometry and n=3 slides from N=1 independent experiment. One-way ANOVA with Dunnett’s correction was used, and data are represented as mean ± standard deviation.

Taken together, these results indicate that FSS does not affect the expression of activation markers or the production of cytokines by naïve BMDMs.

### FSS does not affect the functional responses of classically activated BMDMs

We next examined the effects of FSS on classically activated BMDMs. First, we confirmed that LPS stimulation induces an inflammatory phenotype. Naïve BMDMs were stimulated with 10 or 100 ng/mL LPS for 24 hr, which induced secretion of TNF, IL-6, and IL-10, indicating successful polarisation to an inflammatory phenotype (Supplementary Fig. 1). Naïve BMDMs plated on glass slides were stimulated with 10 ng/mL LPS for 24 hr, after which the supernatant was removed, and LPS-free media was added to the cells. The cells were then subjected to FSS (0.5 or 0.8 Pa, for 10 min/day at 0.5 Hz for two rounds, 24 hr apart), as indicated in Fig. 3A.

**Figure 3:**
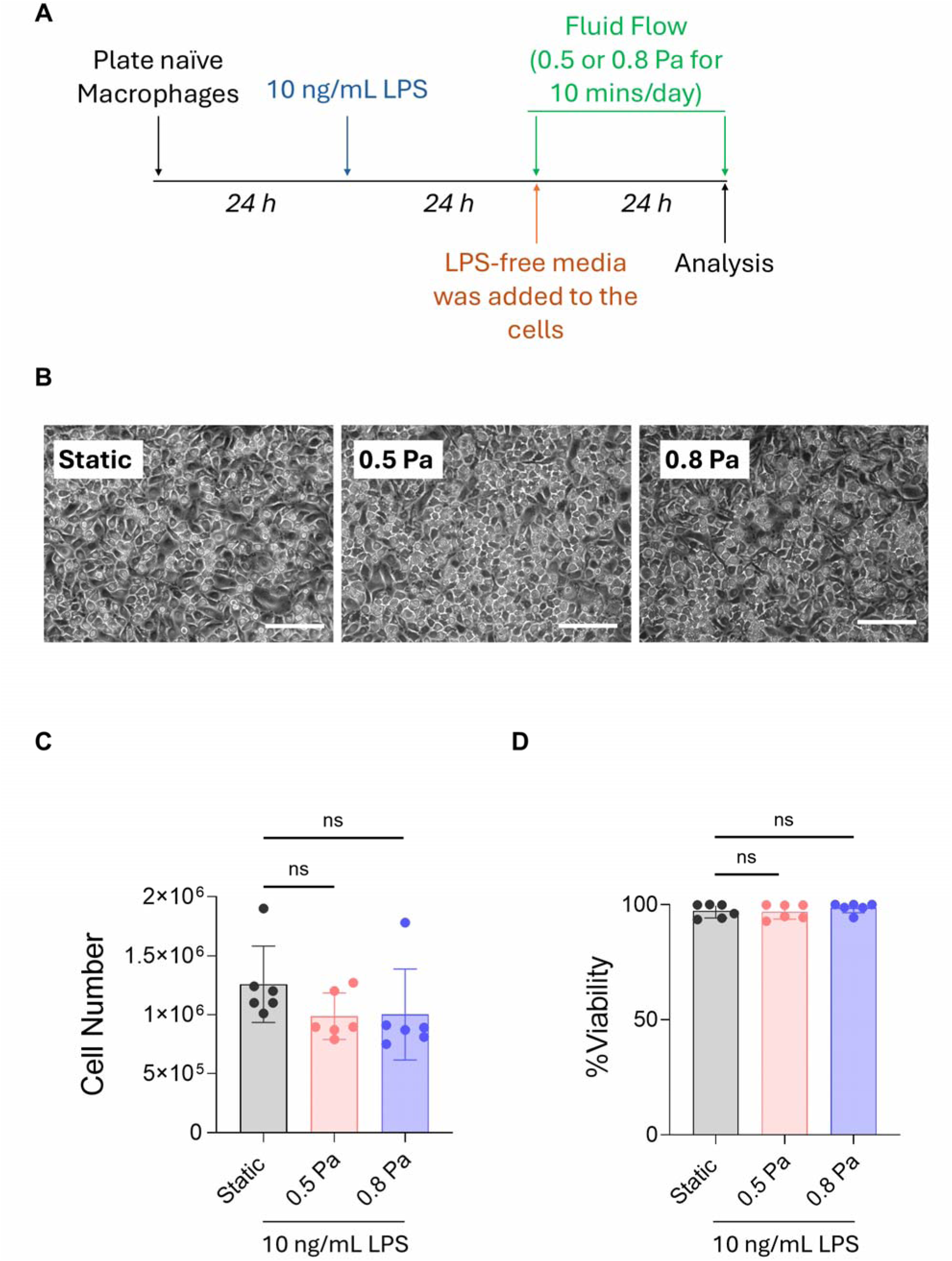
A) Timeline of the study, including 10 ng/mL LPS stimulation for 24 hr prior to FSS exposure (0.5 or 0.8 Pa, for 10 min/day at 0.5 Hz for two rounds, 24 hr apart). B) Microscopic images of the classically activated BMDMs (scale bar: 100 μm), C) cell numbers measured using Luna cell counter and D) percentage of viability in the classically activated BMDMs exposed to FSS measured using Live/Dead Aqua staining and flow cytometry. Data are from n = 6 slides from N = 2 independent experiments. One-way ANOVA with Dunnett’s correction was used, and data are represented as mean ± Standard deviation.

Microscopic images showed that FSS at 0.5 or 0.8 Pa did not affect adherence of classically activated BMDMs to the glass slides and activated BMDMs retained an elongated, spindle-like morphology following FSS at 0.5 or 0.8 Pa (Fig. 3B). There were no differences in the number of cells between the static and FSS groups (Fig. 3C). Furthermore, FSS did not affect the survival of classically activated BMDMs, as cell viability was similar across static and FSS groups (Fig. 3D).

In classically activated BMDMs FSS did not alter the percentage of cells expressing CD11b, CD86 or CD206, or the expression of CD11b and CD86, as measured by MFI, (Fig. 4A-H). However, FSS at 0.8 Pa significantly reduced the expression of CD206, as measured by MFI (Fig. 4I), suggesting that higher magnitudes of FSS may inhibit an anti-inflammatory macrophage phenotype, or impede the resolution of inflammation. We did not observe any differences in TNF, IL-6, IL-10 secretion from LPS-treated BMDMs that were exposed to FSS, as compared to the static control (Fig 4J-L).

**Figure 4:**
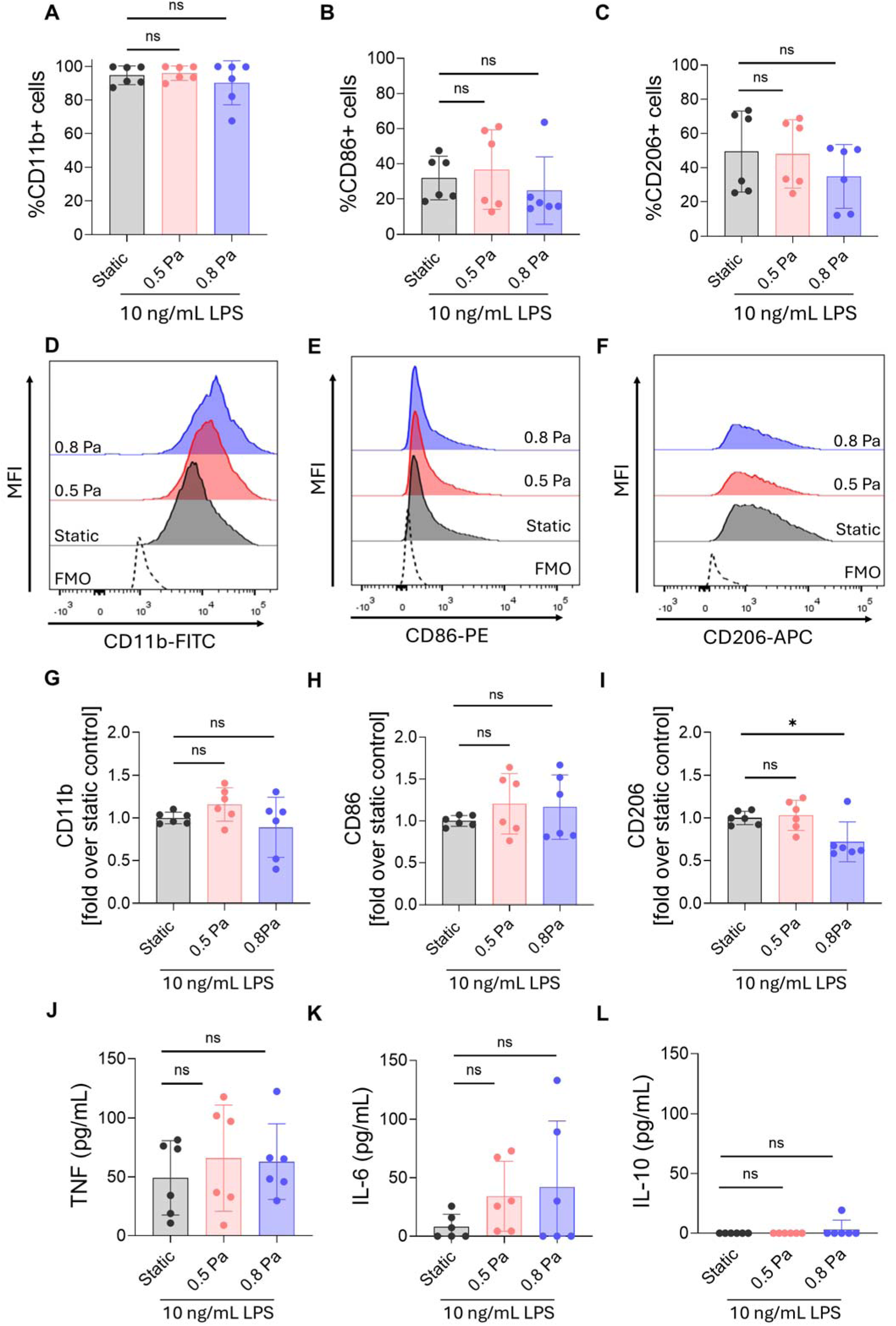
BMDMs were seeded on glass slides and stimulated with 10 ng/mL LPS for 24 h, followed by FSS exposure (0.5 or 0.8 Pa, for 10 min/day at 0.5 Hz for two rounds, 24 hr apart). Percentage of classically activated BMDMs expressing each marker, including CD11b (A), CD86 (B), and CD206(C); representative histograms depict mean fluorescence intensity (MFI) of CD11b, CD86, and CD206 in classically activated BMDMs exposed to FSS (D-F). Pooled data show that FSS does not affect CD11b, CD86, or CD206 expression (G-I), or cytokine production in the supernatant measured by ELISA (J-L). Data are from n = 6 slides from N = 2 independent experiments. One-way ANOVA with Dunnett’s correction was used, and data are represented as mean ± standard deviation.

Taken together, these results indicate that FSS may synergise with inflammatory signals to sustain an inflammatory macrophage phenotype; however, further studies would be required to confirm this response.

### FSS primes BMDMs to drive TNF and limit IL-10 production

Finally, we investigated the effect of exposing BMDMs to FSS prior to LPS stimulation. In this case, naïve BMDMs plated on glass slides were subjected to two rounds of FSS at 0.5 or 0.8 Pa, for 10 mins at 0.5 Hz, once every 24 hr, followed by 10 ng/mL LPS stimulation after the second FSS exposure (Fig. 5A).

**Figure 5:**
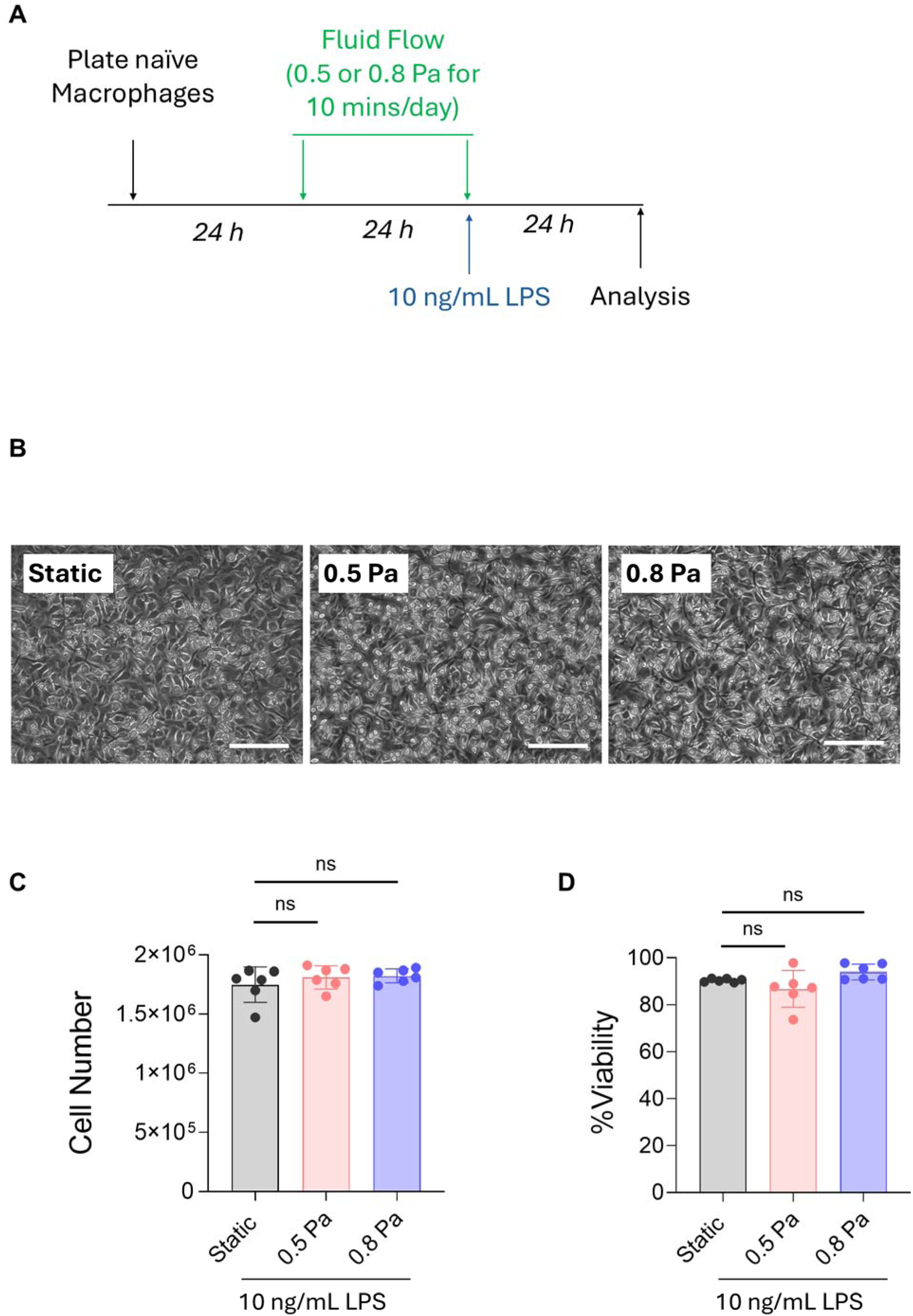
(A) Timeline of the study, including cell seeding on glass slides before FSS exposure (0.5 or 0.8 Pa, for 10 min/day at 0.5 Hz for two rounds, 24 hr apart), followed by 10 ng/mL LPS stimulation for 24 hr, B) Microscopic images of the cells (scale bar: 100 μm), C) cell numbers measured using Luna cell counter and D) percentage of viability in the naïve BMDMs measured using Live/Dead Aqua staining and flow cytometry. Data are from n = 6 slides from N = 2 independent experiments. One-way ANOVA with Dunnett’s correction was used, and data are represented as mean ± Standard deviation.

Microscopic images showed that FSS at 0.5 or 0.8 Pa, prior to LPS stimulation, did not affect cell adherence to the glass slides, and that cells in all groups retained an elongated, spindle-like morphology following FSS at 0.5 or 0.8 Pa and LPS stimulation (Fig. 5B). Furthermore, FSS exposure and LPS stimulation did not affect macrophage survival, as cell viability and numbers were similar across static and FSS groups (Fig. 5C-D).

In this case, FSS did not alter the percentage of cells expressing CD11b, CD86, or CD206, or the expression of CD86 or CD206, as measured by MFI (Fig. 6A-F). However, CD11b expression, as measured by MFI, was significantly increased when cells were exposed to FSS prior to LPS stimulation (Fig. 6G). Alongside this, FSS at 0.8 Pa significantly increased TNF and reduced IL-10 secretion in LPS-activated BMDMs, however there was no difference in IL-6 production (Fig. 6J-L). Of note, cytokine levels of TNF, IL-6, and IL-10 were slightly lower than what has been reported in the literature for LPS stimulation of BMDMs[38,39]. This is likely due to differences in the culture substrate (glass slides vs tissue culture plastic) rather than cell health, as microscopic images confirmed a confluent cell layer with healthy morphology and good adherence across groups, and flow cytometry confirmed good viability.

**Figure 6:**
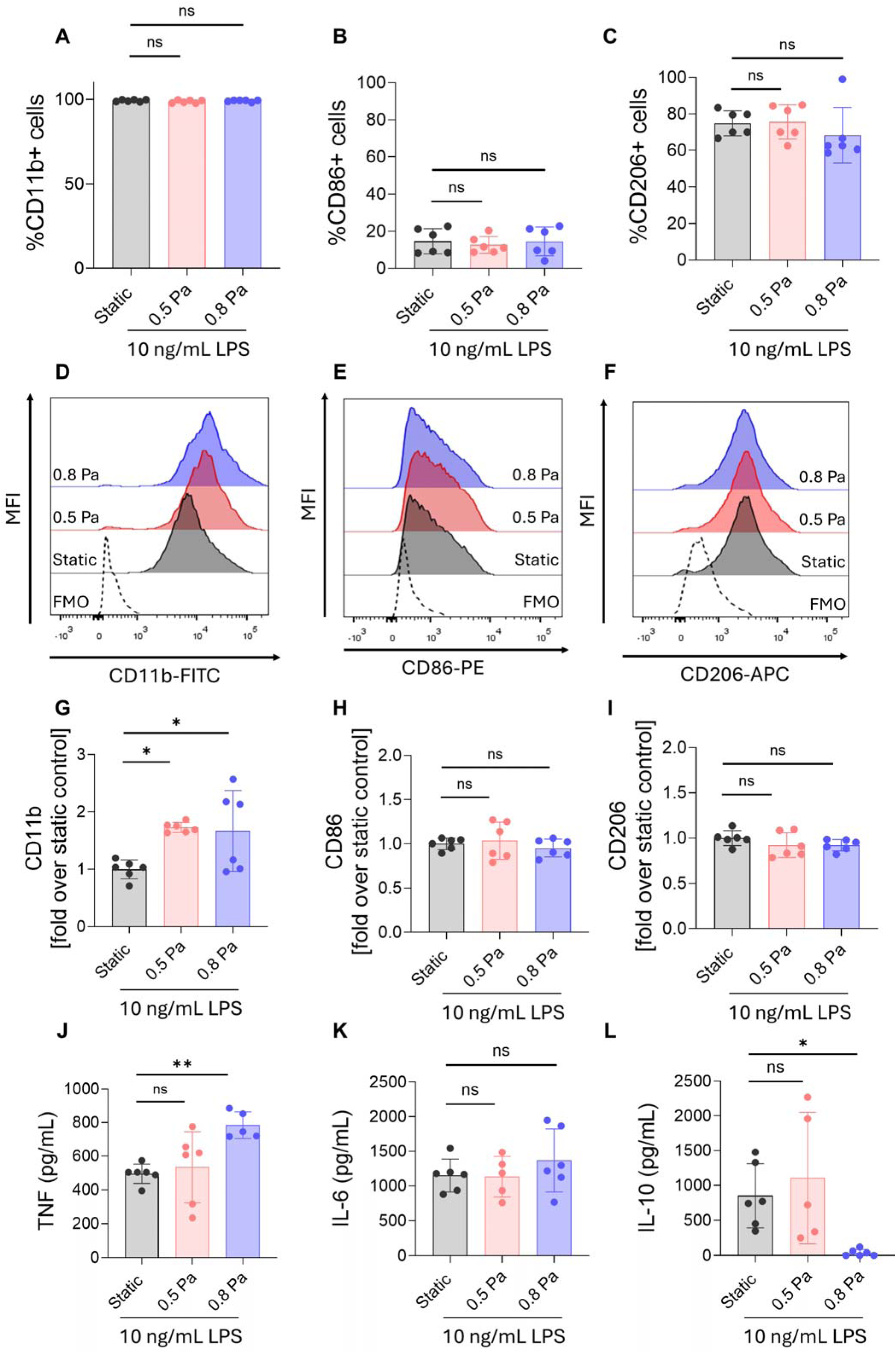
BMDMs were seeded on glass slides before FSS exposure (0.5 or 0.8 Pa, for 10 min/day at 0.5 Hz for two rounds, 24 hr apart), followed by 24 hr of 10 ng/mL LPS stimulation right after the second FSS exposure. Percentage of cells expressing each marker, including CD11b (A), CD86 (B), and CD206(C); representative histograms depict mean fluorescence intensity (MFI) of CD11b, CD86, and CD206 in naïve BMDMs exposed to fluid flow-induced shear stress (D-F). Pooled data showing FSS caused and increase in CD11b fold change, without affecting CD86, or CD206 expression (G-I), and cytokine production in the supernatant measured by ELISA (J-L). Data are from n = 6 slides from N = 2 independent experiments.  One-way ANOVA with Dunnett’s correction was used, and data are represented as mean ± Standard deviation.

Together, these results indicate that short-duration FSS is an inflammatory signal that primes BMDMs to LPS stimulation to support an inflammatory phenotype.

## Discussion

There is a growing recognition that mechanical signals play a key role in shaping immune cell behaviour[40]. Macrophages can sense a variety of mechanical stimuli, including fluid shear stress (FSS), substrate stiffness, pressure and stretch, and translate these signals into changes in inflammatory state[41]. The relationship between FSS and inflammatory state of macrophages is not linear; instead, macrophages appear to exhibit a mechanosensitive threshold response in which the magnitude and duration of applied shear, as well as the original polarisation state of the cells, are critical determinants of the outcome. At lower shear stress magnitudes (approximately 0.01–0.1 Pa, applied over 24–72 hours), several studies have reported a pro-inflammatory shift, characterised by upregulation of TNF, IL-6, and surface expression of inflammation-associated markers[29,42,43]. However, the macrophage response to mechanical stimulation remains highly context-dependent. In particular, the FSS magnitudes and exposure durations used across the literature vary considerably, and many studies employ prolonged or relatively intense loading regimens[17,28,44] that may not reflect the transient mechanical environments encountered by immune cells in the circulation and inflamed tissue environments *in vivo*[45]. Furthermore, it remains unclear whether short mechanical signals are sufficient to directly alter macrophage phenotype or instead modify the subsequent response to inflammatory biochemical cues. To address this, we used a parallel-plate flow chamber to expose bone marrow-derived macrophages (BMDMs) to mild FSS of 0.5 or 0.8 Pa for 10 minutes daily for two days at 0.5 Hz, either alone or in combination with LPS stimulation, with different sequences.

Our findings suggest that mild, short-duration FSS does not act as a strong inflammatory stimulus on naïve BMDMs on its own. Neither 0.5 nor 0.8 Pa significantly affected cell viability, morphology, adherence, surface-marker expression, or cytokine production compared with static controls. However, a different response emerged when the mechanical stimulus preceded biochemical activation. Exposure to 0.8 Pa FSS before LPS stimulation significantly increased TNF production and reduced IL-10 production relative to static LPS-treated BMDMs, whereas 0.5 Pa did not significantly alter the cytokine response. In contrast, applying FSS after 24 hours of LPS stimulation produced no significant changes in TNF, IL-6 or IL-10. These results indicate that short-duration FSS may function as a mechanical priming signal, rather than as a direct activator of resting macrophages, and that this priming effect depends both on the magnitude of the stimulus and on the order in which mechanical and biochemical signals are encountered.

The pro-inflammatory effect observed following a 0.8 Pa pre-exposure is broadly consistent with several studies reporting inflammatory macrophage responses to shear stress. Human monocyte-derived macrophages exposed to sustained laminar FSS have shown increased expression or secretion of TNF, IL-8, IL-18 and other inflammatory mediators, with responses generally increasing with shear magnitude and exposure duration[28]. Similarly, Son et al. reported that 1.2 Pa laminar FSS upregulated CCL2, TNF and IL-1β expression in THP-1-derived cells through heat-shock-protein-associated signalling[43]. Other studies suggest that mechanically induced inflammatory activation can also arise through mechanosensitive ion channels. For instance, it has been demonstrated that FSS primes NLRP3 inflammasome activation through PIEZO1-associated signalling in macrophages and that this response is magnitude-dependent[46]. These observations support the possibility that the increased TNF and reduced IL-10 observed at 0.8 Pa in our study represent a threshold-dependent mechanosensitive response in which FSS alters macrophage responsiveness to subsequent inflammatory stimulation.

Importantly, however, the literature does not support a simple model in which mechanical stimulation universally causes a pro-inflammatory response in macrophages. Interstitial flow has been shown to promote an anti-inflammatory (alternatively activated) response, as evidenced by upregulation of ARG1, TGF-β, CD206, CD163, and TGM2 via β1-integrin/Src-dependent JAK–STAT signalling[11]. Likewise, different mechanical modes can suppress inflammatory signalling; for example, cyclic stretch has been reported to inhibit ATP-induced IL-1β secretion in LPS-primed macrophages by inhibiting NLRP3 inflammasome activation[47]. These apparently contradictory findings are likely to reflect fundamental differences in the mechanical environment rather than genuine disagreement between findings. Shear magnitude, duration, cell culture system design, macrophage cell source and pre-existing polarisation state can all determine how a mechanical signal is interpreted. The macrophage response to FSS should therefore be considered a spectrum rather than a binary pro-versus anti-inflammatory state.

The present study contributes to this framework by highlighting the importance of both magnitude, exposure duration and sequence of stimuli. The absence of a measurable response at 0.5 Pa, combined with the increased TNF and reduced IL-10 observed at 0.8 Pa, suggests that a threshold may exist between these magnitudes under the conditions tested (taking into account the frequency and duration of exposure). Moreover, the observation that the response was observed when FSS was applied before LPS, but not when FSS was applied after LPS stimulation, suggests that macrophage mechanosensitivity depends strongly on the cell state at the time the stimulus is applied. This is consistent with broader evidence that biochemical and mechanical signalling pathways interact rather than operate independently. PIEZO1, for example, has been implicated not only in mechanical sensing but also in inflammatory macrophage polarisation and in interactions with TLR4 signalling[48]. Given that LPS is a strong stimulus that elicits robust TLR4-dependent activation even at very low concentrations[49]; the LPS-induced response may have overshadowed subtler FSS-specific effects in this system. Thus, a brief mechanical signal may alter intracellular signalling or cellular readiness without producing a large immediate inflammatory response, subsequently changing the magnitude of the response when a biochemical stimulus is encountered.

This is particularly relevant because relatively little attention has been given to short-duration FSS exposures. Much of the existing macrophage literature has used shear exposure lasting from tens of minutes to several hours or even days. For example, studies included in the literature comparison applied shear for 30 minutes or 5 hours, applied prolonged laminar shear at multiple time points, or applied repeated mechanical loading for 1 hour per day over several days[28,50]. By contrast, the 10-minute regimen used here was designed to investigate whether a brief mechanical exposure could influence macrophage behaviour without reproducing the longer sustained loading conditions commonly studied. Such transient exposure is more representative of the mechanical environment encountered by the immune cells *in vivo*, where monocyte contact with the endothelium under flow lasts only seconds[51]. The lack of direct activation following this short exposure, coupled with altered responsiveness to subsequent LPS, indicates that the biological consequences of transient FSS may be more subtle than those produced by prolonged loading and may manifest primarily through altered sensitivity to later inflammatory signals.

The findings of this study are particularly relevant to our group’s ongoing work on actuatable medical implants. Previous studies from our group have demonstrated that short-duration mechanical actuation of an implant can perturb cellular activity at the implant interface to modulate the foreign body response and improve drug delivery through the actuatable implants[52,53]. We showed that actuation of the implant (10 mins/day) significantly reduced the presence of neutrophils at day 5[52] and myofibroblasts at day 14[52,53] in the peri-implant tissue. Interestingly, and of particular relevance to the current study, actuation did not influence macrophage number at the implant surface at day 14[53]; however, macrophage phenotype was not characterised, and day 14 was potentially too late to capture the acute macrophage response. Computational modelling predicts that actuation generates local interstitial fluid flow (<50 mm/s[52], roughly equates to 1 Pa) and tissue strain (up to 25%[52]), meaning that macrophages adjacent to the implant are repeatedly exposed to combinations of mechanical stimuli, in addition to biochemical signals. The current study isolates the FSS component of this environment and demonstrates that even short mechanical exposure (10 mins/day) can modify subsequent macrophage inflammatory behaviour *in vitro*. Notably, cell number was maintained across FSS conditions, consistent with the *in vivo* finding by Dolan et al.[53], which reported no significant changes in numerical density of macrophages at the implant interface following actuation. The increased TNF and reduced IL-10 observed following 0.8 Pa FSS should therefore not necessarily be interpreted as inconsistent with the reduced fibrosis observed following longer-term implant actuation. The foreign body response is temporally dynamic, progressing through acute inflammation, resolution, and ultimately fibrotic remodelling[54]. A short-term increase in inflammatory responsiveness may contribute to early host defence, recruitment and clearance processes, while repeated mechanical stimulation over subsequent days may have different effects as macrophages transition between inflammatory states. Mechanical cues may therefore influence different stages of the foreign body response in distinct ways rather than maintaining a uniformly pro- or anti-inflammatory effect over time. Determining whether repeated actuation ultimately redirects macrophages toward a more classically or alternatively activated state will require longitudinal investigation of macrophage phenotype and cytokine production at multiple stages following implantation *in vivo*.

## Limitations of this study

There are some limitations to our study. First, macrophages cultured on rigid glass substrates experience FSS in an environment that does not recapitulate the compliant, heterogeneous extracellular matrix present *in vivo*. In tissues, macrophages are exposed simultaneously to shear, strain, compression, substrate stiffness, and biochemical signals, and these inputs may interact in ways that cannot be reproduced in a reductionist parallel-plate system. Second, only two mild FSS magnitudes were examined, limiting the ability to define the precise mechanical threshold or determine how repeated exposure changes the response. Finally, only selected cytokines and surface markers were assessed, and additional studies will be required to determine whether pathways such as PIEZO1, integrin signalling or other mechanosensitive mechanisms mediate the priming response observed here. Nevertheless, the system used in this study provides a controlled means of isolating FSS and demonstrates that macrophage responses to FSS depend not only on the presence of mechanical stimulation, but also on its magnitude, duration and sequence of mechanical and biochemical stimulation.

## Conclusion

Our results indicate that mild FSS alone was insufficient to activate naïve BMDMs, whereas a brief 0.8 Pa exposure before LPS enhanced TNF production and suppressed IL-10. The absence of this response at 0.5 Pa and when shear was applied after LPS indicates that both a mechanical threshold and the stimulation sequence are important determinants of macrophage behaviour. Taken together, these findings support a model in which transient mechanical exposure serves as a priming signal that tunes macrophage responses to subsequent inflammatory stimuli, consistent with existing evidence that shear stress can act as a discrete signal in macrophage activation pathways, depending on stimulus sequence and magnitude. This provides a potential mechanism through which mechanically active implants and other dynamic tissue environments regulate innate immune function.

## Acknowledgements

E.B.D., P.S., and H.P. acknowledge funding from the Research Ireland Starting Laureate Award (IRCLA/2022/2766). E.B.D., P.S., H.P. and R.D. acknowledge funding from the Research Ireland Royal Society University Research Fellowship (URF\R1\191335, URF\ERE\210376, URF\R\241016).

E.B.D and H.P. acknowledge funding from the Government of Ireland Postdoctoral Fellowship (GOIPD/2024/740). H.P. acknowledges Research Seed Funding from the College of Science and Engineering at University of Galway (AY23/24).

The authors thank Dr. Juan Alberto Panadero Pérez and Dr. Vatsal Kumar for technical support, and Dr. Shirley Hanley for assistance with flow cytometry. The authors also thank Prof. Laoise McNamara, Biomedical Engineering at the University of Galway, for providing their custom-made parallel-plate flow chambers.

## Authors Contributions

E.B.D., H.P. and P.S. conceived and designed the study. P.S. performed the experiments, collected and analysed the data, and wrote the original draft of the manuscript. R.D. contributed to the experimental setup. E.B.D. and H.P. provided study materials and contributed to data analysis and interpretation. E.B.D. and H.P. supervised the project and coordinated the work. All authors contributed to the writing, critically reviewed, and approved the final version of the manuscript.

## Supplementary Data

**Supplementary Figure 1:** A) Timeline of the experiment, where BMDMs were seeded on glass slides and rested for 24 h prior to LPS stimulation for 24 hr. Production of B) TNF, C) IL-6, and D) IL-10 by BMDMs stimulated with 0, 10 or 100 ng/mL LPS.

**Supplementary Table 1:** Antibodies used for flow cytometry.

| Antibody | Clone | CAT Number | Dilution | Supplier |
|---|---|---|---|---|
| CD11b | M1/70 | 101205 | 1:200 | BioLegend |
| F4/80 | BM8 | 123113 | 1:200 | BioLegend |
| CD86 | A17199A | 159203 | 1:200 | BioLegend |
| CD206 | C068C2 | 141707 | 1:200 | BioLegend |

## Resource Availability

### Lead Contact

Requests for further information and resources should be directed to and will be fulfilled by the lead contact, Hannah Prendeville.

### Materials Availability

This study did not generate new unique reagents. All antibodies used in this study are commercially available and listed in the Supplementary Table (Table S1).

### Data and Code Availability

All raw data reported in this paper will be shared by the lead contact upon request. This study did not generate new custom code. Any additional information required to reanalyse the data reported in this paper is available from the lead contact upon request.

