## Supplementary figures and images for "Short-Duration Fluid Shear Stress Enhances LPS-Driven Macrophage Pro-Inflammatory Response in a Magnitude-Dependent Manner"

### Supplemetary Figure 1

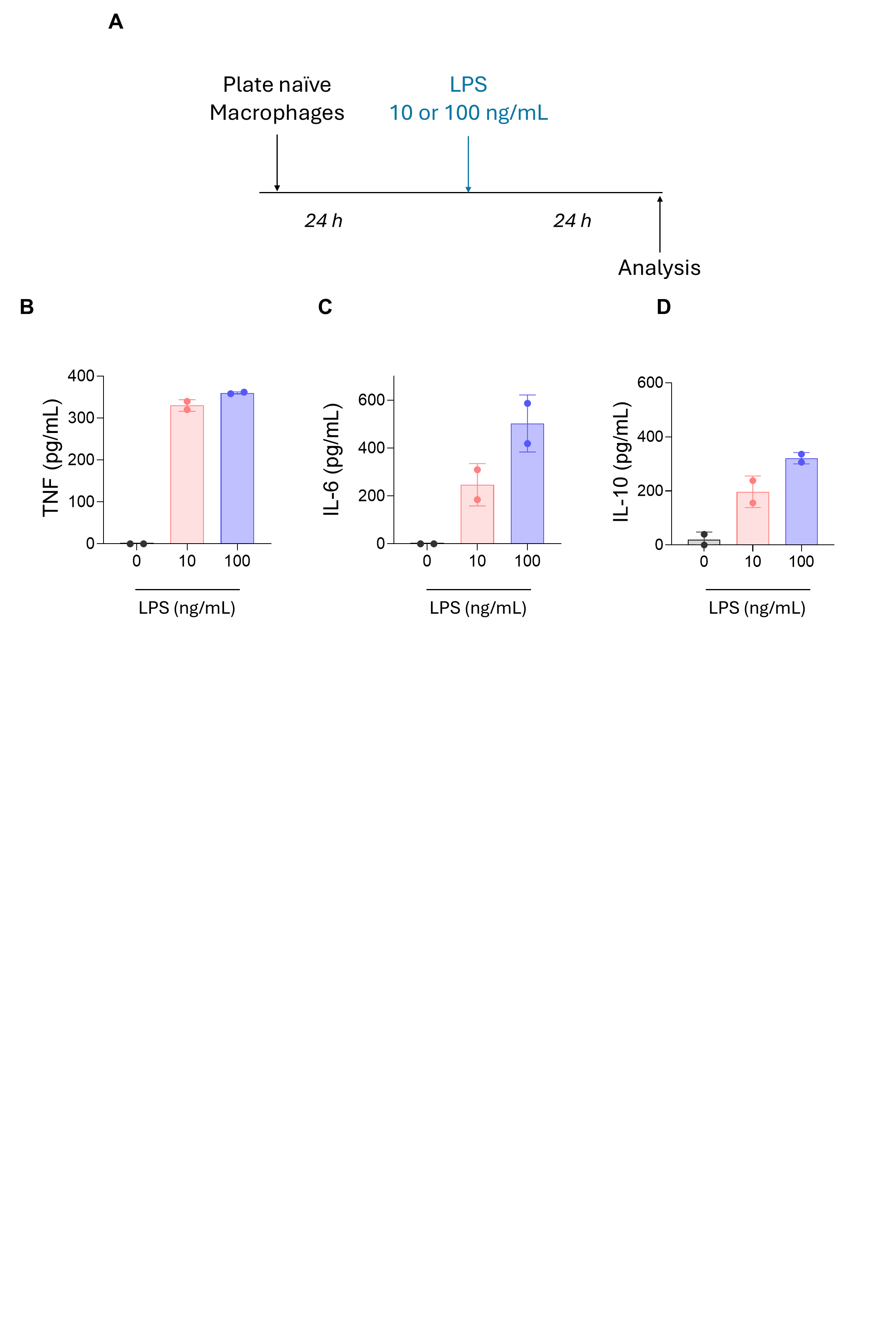
